# A Window into the Brain’s Microcirculation: Retinal Optical Coherence Tomography Angiography (OCTA) as Perioperative Bedside Resuscitation Guide?

**DOI:** 10.1101/2025.03.03.640442

**Authors:** Sourav S. Patnaik, Irtiza Sakif Islam, Jai Singh Rajput, Kasandra Albarran, Aksharkumar Dobariya, Misha Dunbar, Juan M Pascual, Ulrike Hoffmann

## Abstract

**Objective:** Quantifying physiological flow dynamics in the brain’s microvasculature is vital for effective resuscitation management. Traditional resuscitation approaches rely on macrocirculatory flow targets, which often poorly correlate with cerebral microcirculatory flow. Our observational pilot study investigates the feasibility of using optical coherence tomography angiography (OCTA) to image retinal microcirculatory blood flow in the immediate perioperative setting.

**Methods:** Using a porcine animal model, we were able to induce *hypercarbia*, epinephrine-led *resuscitation,* and hemorrhage followed by autologous *re-transfusion* scenarios.

**Results:** Vascular density of superficial capillary plexus showed an average increase in perfusion of approximately 4% and 1.2% from baseline for *resuscitation* and *hypercarbia* stages, respectively. Conversely, *re-transfusion* stage showed an approximate average reduction in superficial layer vascular density by 3.9% from baseline. Vascular density of the deep capillary plexus layer showed a significant increase (6.31%) from baseline in for *hypercarbia* stage but remained unchanged for the rest of the stages.

**Conclusions:** Taken together, OCTA can be (i) utilized in a perioperative setting, (ii) used to detect fast changes in systemic blood pressure, and (iii) utilized for non-invasive cerebral blood flow pattern determination during pre- and post-surgical evaluation of patients.

**Disclosures:** None

## Introduction

Impairment of microcirculatory blood flow has been implicated as a pivotal pathophysiologic event perioperatively and in critically ill patients. As the brain is particularly sensitive to insufficient perfusion, resulting cognitive and neurologic impairment can prolong the recovery period and significantly impact the ability to live independently.

To date, resuscitation or perfusion management in patients is primarily guided by macrocirculatory assessment (i.e., arterial blood pressure) with little consideration of the microcirculation, as microcirculatory parameters are technically difficult to assess. However, recent studies in pathophysiologic states (e.g., sepsis, shock) strongly suggest that microvascular perfusion is not restored despite the optimization of macrocirculatory parameters. Thus, there is an urgent need to deduce what governs cerebral microcirculatory flow and define effective microcirculation resuscitation targets.

While research tools exist, there is no commonly accepted technique which can measure microcirculation through the intact skull. Therefore, it is essential to identify more accessible microcirculatory beds (e.g., retinal) which may serve as a window into and surrogate for cerebral microcirculatory flow. Characterizing surrogate microcirculations as flow-biomarkers is a critical step towards practical clinical implementation of cerebral microcirculatory targets for resuscitation. In particular, the retinal microcirculation can be considered as window into the cerebral microcirculation, as the retina and brain share similar microvascular anatomy, and retinal structural and blood flow changes associated with systemic and central nervous system illness are increasingly reported [1–4]. Retinal and cerebral microvasculature have a common blood supply via the internal carotid artery which supplies blood to the ophthalmic artery and central retinal artery. Both cerebral and ocular microcirculatory beds have similar physiology in healthy and diseased conditions. A recent OCTA study of 137 clinical patients with no known neurological diseases showed that reduced retinal perfusion is associated with reduced gray matter volumes in brain regions (putamen, occipital, frontal, and temporal lobes), and positively associated with structural covariance of major parts of the brain [5]. Retinal microcirculation changes may, therefore, correlate with cerebral microcirculatory dynamics in critically ill patients, offering a novel biomarker (a *window* into the brain) to monitor in real time at bedside [4].

Optical coherence tomography angiography (OCTA) is currently predominately used for diagnosing ocular diseases; here we propose OCTA as a non-invasive tool to detect retinal microcirculatory dynamics in tandem with systemic blood pressure (BP) changes. This novel technology provides high-resolution images of retinal and choroidal vascularization and permits *in vivo* measurement of microcirculatory blood flow dynamics inaccessible in any other microcirculation. This technology does not need dyes or contrast agents, is portable, non-invasive, reproducible, and, with some future biomedical modification, could be performed at bedside. In consequence, OCTA may be suitable for assessing dynamic functional measures of the retinal microcirculation (e.g., flowmetry and dynamic vessel assessments).

In this study we investigated OCTAs ability to capture acute, fast changing physiological scenarios and to quantify vascular flow features of the retinal superficial capillary plexus layer (SCP) and deep capillary plexus layer (DCP) in a porcine model under various physiological challenges to determine its utility to function as a future online feedback parameter for clinical scenarios such as hemorrhage and resuscitation, to guide clinical management.

## Materials and Methods

### Animal preparation

All animal experiments were conducted in accordance with protocols approved by the University of Texas Southwestern Institutional Animal Care and Use Committee (IACUC) and in accordance with ARRIVE guidelines (Animal Research: Reporting of In Vivo Experiments). Juvenile farm animals (*Sus scrofa domesticus,* Yorkshire cross) about 2– 4-month-old (one male and one female, ∼20-40 kgs) were procured 1-2 weeks prior to the experiment and acclimatized in the Association for Assessment and Accreditation of Laboratory Animal Care International accredited UT Southwestern facility (elevated pens on tenderfoot or slatted flooring). Animals were fed with routine diet (Teklad Mini swine Diet 7037, Envigo, Madison, WI) and ad-libitum water. Standard housing temperature (18-22°C) and humidity control (30-70%) were maintained with a 12-hour light/dark cycle.

### Anesthesia and physiological monitoring

Thirty minutes prior to surgery, animals were anesthetized using routine cocktails of anesthetics and analgesics (telazol 4-8mg/kg; glycopyrrolate 0.005 mg/kg; ketamine 10-20mg/kg; xylazine 2mg/kg) via intramuscular route, per previously established protocols [6, 7]. Using a facemask, 1.5-2% isoflurane and oxygen (2L/min) was administered to the animals were intubated. An A-line catheter was placed on the left femoral artery for arterial blood pressure monitoring and blood gas retrieval as well as a femoral intravenous catheter for blood removal. Animals were placed in prone position on the operating table, esophageal EKG probe and rectal temperature probes were placed for physiological monitoring. A mechanically assisted respirator was utilized to maintain ventilation at 15-20 breaths per minute. Normal saline (Normasol-R, ICU Medical, Inc., San Clemente, California) was administered to the animals throughout the experiment at the rate of 200-250ml per hour. A strap was placed around the torso to stabilize the body during the experiment and a heated air blanket (Bair hugger, 3M, MN) was utilized to maintain body temperature. Animal vital signs (arterial BP, end-tidal CO_2_, oxygen saturation, etc.) (SurgiVet Advisor® tech Vital Signs Monitor, Smiths Medical, Inc., Minneapolis, MN, AD instruments) were recorded throughout the experiment.

### Experimental Stages

The overall layout of the study is shown in Figure 1 and detailed experimental stages are provided in Table 1. The primary objective of this study was to capture microvascular flow changes in the porcine retinal capillary network (SCP and DCP layers) using OCTA imaging in parallel to systemic hemodynamic changes in the animal. We investigated the ability of OCTA to capture fast changing, clinically relevant physiological conditions: (i) *baseline* (anesthetized condition); (ii) *hypercarbia* (target PaCO_2_ of 70 mmHg, confirmed by blood gas measurements); (iii) *hypotension* by increased dose of isoflurane, (iv) *resuscitation* using epinephrine bolus, (v) blood withdrawal (target systemic BP 40-45mmHg; about 30% of total blood volume; 30 mL/kg representing ∼800 mL within 30 minutes [9]); followed by (vi) autologous *re-transfusion*. Animals were allowed to recover to normoxic baseline physiological conditions after hypercarbia or normotonic baseline after one epinephrine bolus. We targeted a stability of 90% of baseline systolic blood pressure as the minimum requirement for resuscitation of hypotension [10]. Systolic pressure (SP), diastolic pressure (DP), mean arterial pressure (MAP), oxygen saturation (SpO_2_), end-tidal CO_2_ (EtCO_2_), and heart rate (HR) were recorded for all the experimental stages. At the conclusion of the study, euthanasia was performed under general anesthesia with an intravenous Euthasol bolus (Virbac Corporation, Westlake, TX).

**Fig. 1.**
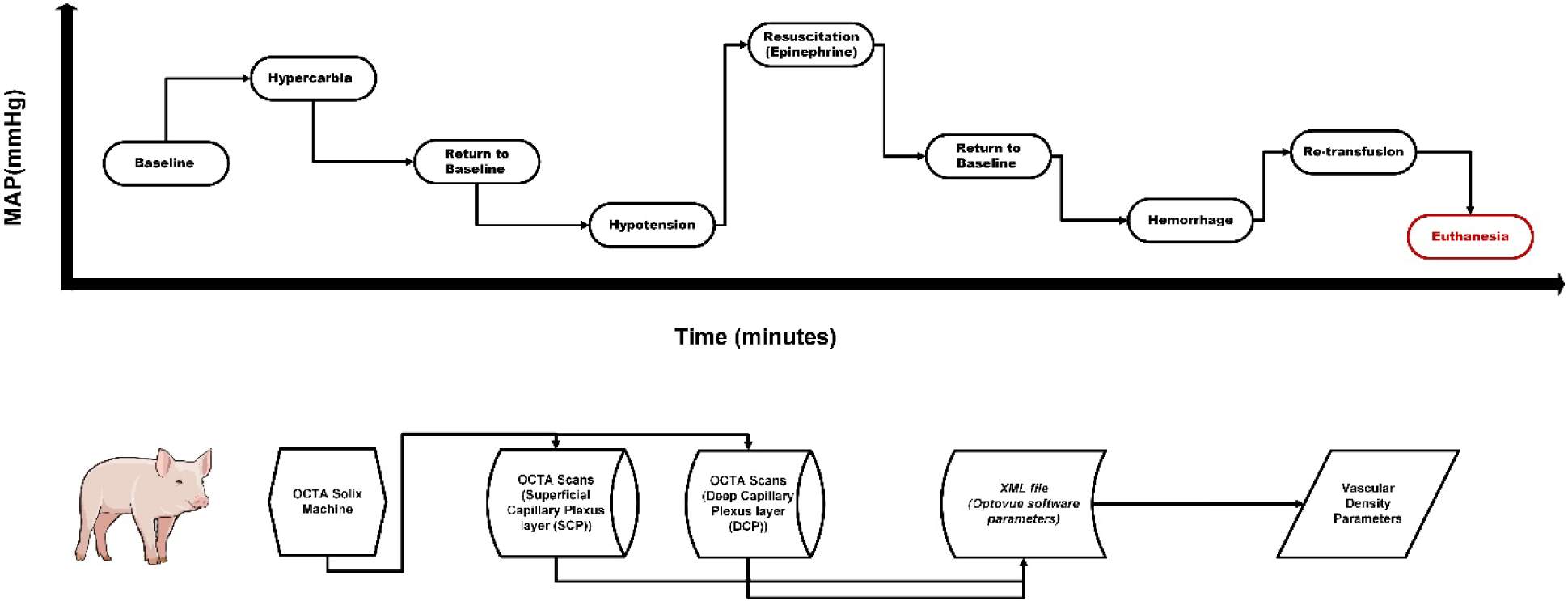
Overall layout of the experimental OCTA study with porcine animals undergoing physiological challenges. Porcine figure courtesy NIAID [8].

**Table 1.** Experimental stages of the study with detailed interventions are listed here.

### Optical coherence tomography angiography protocol

#### OCTA Imaging

The left eye of the animal was utilized for capturing OCTA images. A speculum was used to keep the eye assessable. Frequent saline flushes were performed to keep the eyes moisturized. OCTA imaging of the porcine eyes were performed using Optovue ® Solix acquisition system (Visionix International SAS (formerly Optovue), Pont-de-l’Arche – France) (15 µm/pixel sampling density [11], light source of 840nm; scan speed 120kHz; axial resolution 5µm; lateral resolution 15µm; scan depth up to 3mm; scan width 3-16mm). The head of the animal was manually positioned close to image capture unit, and for each acquisition the optimal focal length (prompted by acquisition software) had to be achieved in order to capture the OCTA scan. The animals were draped with black cloth to facilitate hassle-free OCTA acquisitions. For each experimental stage, multiple OCTA scans of the porcine macular region were performed using the *AngioVue Retina* mode (6.4mmx6.4mm scans, 512×512 pixel resolution). Automatic layer specific segmentation of retinal microvasculature was performed in the *AngioVue* software. Acquired OCTA scans exhibiting motion artifacts or signal quality index of less than 6 were not utilized for further analysis [12].

#### OCTA Analysis

We focused our analysis on the superficial capillary plexus (SCP), defined by retinal anatomical structure from inner limiting membrane to inner plexiform layer, and deep capillary plexus (DCP), defined as retinal anatomical structure from outer boundary of inner plexiform layer to the outer plexiform layer [13, 14]. The primary outcome of this study was vascular density of whole image (VD), superior (VD_Hemi-S_) and inferior (VD_Hemi-S_) hemi-regions of the retinal layers (SCP or DCP) [15]. Signal quality metric was also analyzed across the experimental stages.

### Statistical Analysis

Test for normality was performed using Shapiro–Wilk test. All parameters were compared across the experimental stages using either ANOVA or Kruskal-Wallis test and pairwise comparison as well (Tukey’s or Dunn’s – both tests account for family-wise error rate or false discovery rate). To further explore the relationship between vascular density (both SCP and DCP layers) and experimental parameters, we performed Pearson or Spearman correlation analysis. All analyses were performed in GraphPad Prism and data was considered significant at *p*<0.05.

## Results

Experimental acquisition of OCTA images with the current machine <u>in porcine</u> animals was very time consuming; some of the changes in systemic vascular physiological conditions and its corresponding change in retinal vascular flow, proved to be impossible capturing with high quality OCTA acquisition. In particular, the severe *hypotension* or deep anesthesia stage resulting in relevant reduction of blood pressure and relaxation of retinal blood vessels proved challenging and only very few or no OCTA flow signals were captured. Similarly, the OCTA scans captured during the maximum *hemorrhage* stage (which involved removal of ∼800 ml of whole blood porcine) were not free of error or artifacts, and hence, no good quality data that could be utilized for statistical analysis were available for these two stages.

Henceforth, the experimental stages of this study will refer to *baseline*, *hypercarbia*, *resuscitation* by epinephrine, and the result of autologous *re-transfusion*.

### Systemic parameters

Overall, systemic parameters i.e., arterial systolic pressure (SP), diastolic pressure (DP), mean arterial pressure (MAP), heart rate (HR) and SpO_2_ showed the expected and well-known responses to the physiologic challenges facilitated in our experiments. Baseline values in the respective parameters did not differ significantly from pig to pig, or between female and male gender. During hemorrhage, blood pressure was initially compensated and started to rapidly decrease to the nadir at maximum blood withdrawal (40mmHg) concurrent with a steep compensatory increase in heart rate. One dose of epinephrine was administered, which resulted in the immediate increase of both blood pressure parameters and heart rate as depicted in Figure 2B-D. SP was highest for the epinephrine *resuscitation* stage (126.0±37.07 mmHg), followed by *hypercarbia* (75.5±5.29 mmHg)*, baseline* (63.57±8.52 mmHg) and *re-transfusion* (54.88±5.78 mmHg) stages (*p*<0.0001; Fig.2A). Similar pattern was exhibited by DP(64.25±19.82 mmHg vs. 44.63±7.19 mmHg vs. 37.14±7.56 mmHg vs. 30.56±4.68 mmHg; *p=0.0003*), and MAP (84.83±24.68 mmHg vs. 54.92±5.88 mmHg vs. 45.95±7.57 mmHg vs. 38.67±4.45 mmHg; *p*<0.0001) (Fig. 2B-C), respectively. As expected, HR was higher for epinephrine *resuscitation* stage, followed by the *baseline* being slightly (non-significantly) higher than *hypercarbia* and *re-transfusion* stages (181±15.19 BPM vs. 97.86±14.10 BPM vs. 90±9.64 BPM vs. 86.94±11.39 BPM; *p=0.0045*) (Fig. 2D), respectively.

**Fig. 2.**
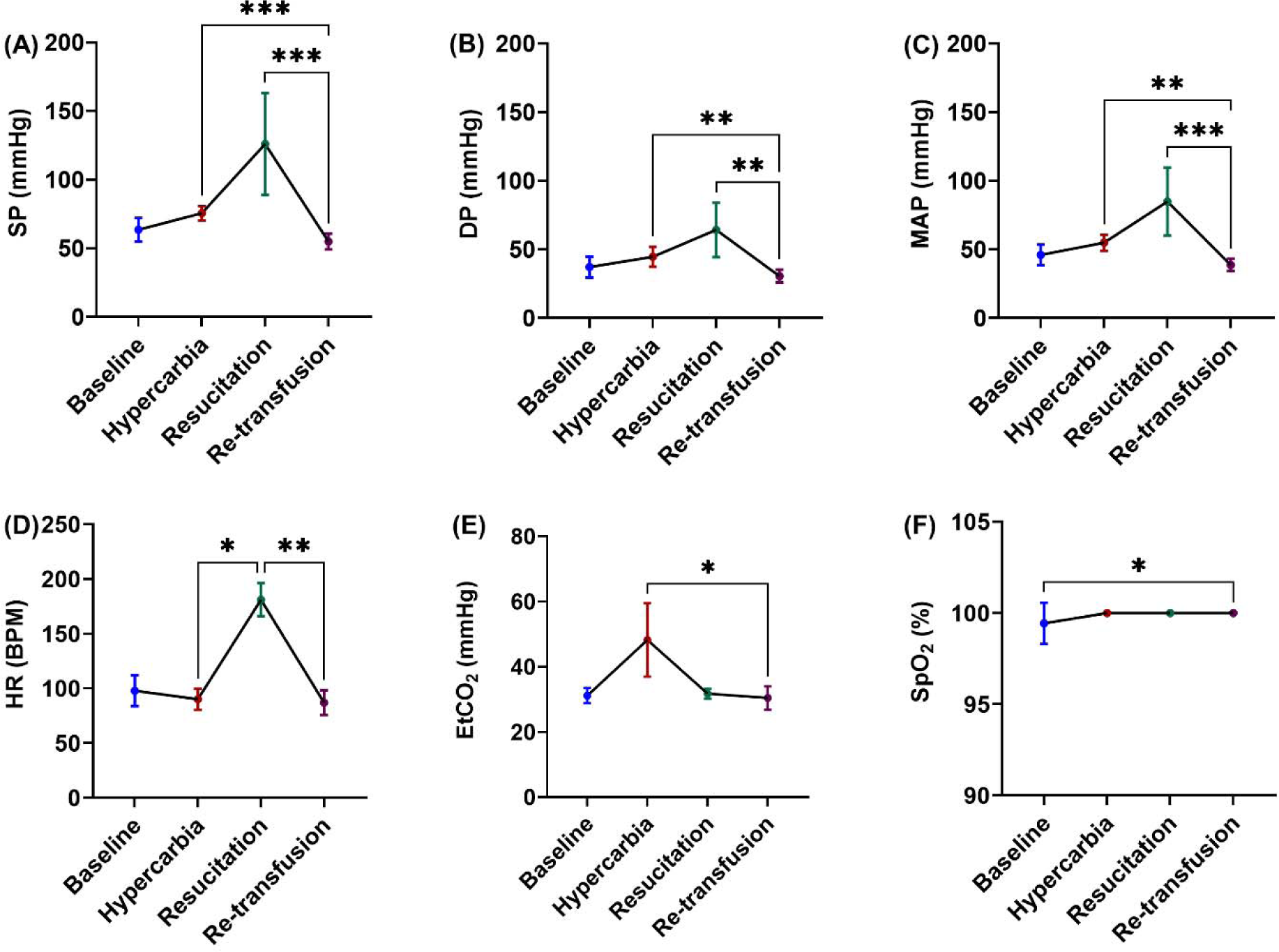
Experimental parameters (systolic pressure (SP) (A), diastolic pressure (DP) (B), mean arterial pressure (MAP) (C), heart rate (HR) (D), end-tidal carbon dioxide (EtCO_2_) (E), and oxygen saturation (SpO_2_) (F)) across baseline, hypercarbia, resuscitation, and re-transfusion stages are reported here.

As expected, EtCO_2_ was highest for *hypercarbia* as compared to the other stages (48.25±11.27 mmHg (*hypercarbia*) vs. 31.14±2.34 mmHg (*baseline*) vs. 31.75±1.5 mmHg (*resuscitation*) vs. 30.44±3.58 mmHg (*re-transfusion*) (*p*=0.0203); Fig. 2E) as was confirmed by a reduction of pH and corresponding increase in PCO_2_ concentrations in blood gases (detailed of blood gas analysis provided in S1 supplementary information).

On the other hand, SpO_2_ was fairly consistent throughout the experiment with minimal changes from *baseline* throughout the end of the experiment after *re-transfusion* of all autologous blood (99.43±1.13% vs. 100% (*p*=0.0414); Fig. 2F), whereas PaO_2_ showed a reduction during hypercarbia, which resolved as CO_2_ levels were normalized.

### OCTA Vascular Density Metrics

In concordance with previously published literature [16–20], no foveal avascular zone was observed in porcine retina from initial gross observation of OCTA images. Signal quality of the OCTA images (scored automatically out of 10) reduced progressively throughout the study (*baseline*: 8.3 ± 1.25 vs. *hypercarbia*: 7.08 ± 1.12 vs. *resuscitation*: 8.0 ± 0.71 vs. *re-transfusion:* 7.05 ± 0.39; *p* = 0.0066).

Fig. 3 A-H Shows representative changes in vascular density across the physiological conditions. We observed that the vascular density of the superficial capillary plexus was highly correlated with systemic physiological changes, whereas the deep capillary plexus displayed seemingly independent dynamics. Gross observations reveal some of the microvasculature appear black? in the *resuscitation* or *re-transfusion* phases as systemic blood flow is restored in the animal.

**Fig. 3.**
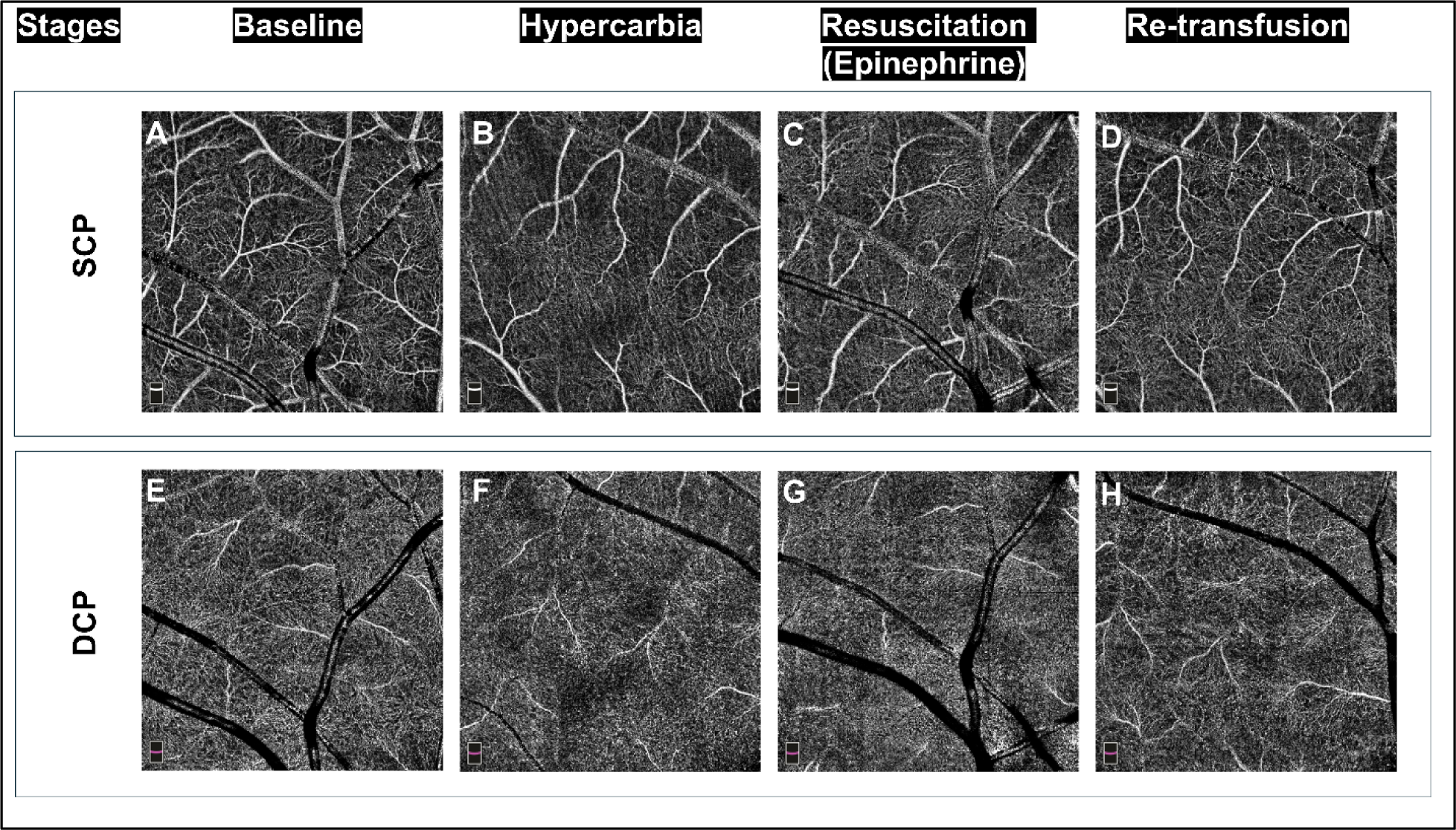
Exemplary porcine OCTA scans (superficial capillary plexus (SCP) (A-D) and deep capillary plexus (DCP) layers) (E-H) captured across baseline, hypercarbia, resuscitation, and re-transfusion stages are shown here.

### Superficial capillary plexus *(SCP)*

SCP vascular perfusion slightly increased during hypercarbia from baseline (*baseline*: 46.65 ± 1.74% vs. *hypercarbia*: 47.19 ± 1.56%) and increased further in response to one dose of epinephrine (resuscitation) paralleling the increase in systemic BP (*baseline*: 46.65 ± 1.74% vs *resuscitation*: 48.50 ± 2.04%). After hemorrhage and full autologous re-transfusion however, vascular density was diminished and did not recover to baseline *(baseline*: 46.65 ± 1.74% vs. *hemorrhage and re-transfusion:* 44.83 ± 1.90%; *p* = 0.0007) (Fig.1B) The drop in macular perfusion was more prominent in the inferior region (Fig.1D) than in the superior region (Fig.1C) as compared to baseline.

### Deep capillary plexus (DCP)

Vascular perfusion in the DCP increased during hypercarbia and remained elevated through the rest of the experiment. We did not observe a change in perfusion in response to epinephrine (*baseline*: 48.46 ± 0.70 % vs. *hypercarbia*: 51.52 ± 1.58% vs. epinephrine *resuscitation*: 50.34 ± 0.99% vs. *full re-transfusion after hemorrhage:* 50.58 ± 1.48%; *p* = 0.0003) (Fig.2B). The superior DCP region showed increased vascular density (Fig.2C), while no such change was observed in the inferior region (Fig.2D).

Spearman correlation analysis did not show any association between VD-SCP and experimental parameters (SP, DP, MAP, HR, EtCO_2_ and SpO_2_) (Table 2; p>0.05). Conversely, VD-SCP exhibited significant, positive associations with SP (*r*=0.4; *p=*0.0176), DP (*r*=0.44; *p*=0.0081), MAP (*r*=0.43; *p*=0.0096), and HR (*r*=0.39; *p*=0.0207). No associations were found between VD-SCP and EtCO_2_ or SpO_2_ (*p*>0.05 for both).

**Table 2.** Correlation of vascular density (SCP and DCP) with physiological parameters. Bold text shows significant correlations between variables (p<0.05).

## Discussion

In this unique investigation, we accomplished three-fold objectives – (i) retinal OCTA imaging is capable of assessing acute physiological changes in real time (ii) retinal flow changes occur in tandem with systemic hemodynamics changes and (iii) the porcine model has similar ocular features to human and can be used to study brain-retina microcirculatory flow.

We observed that the superficial capillary layer (Fig.3A-D) is more sensitive to systemic hemodynamic changes, such as blood withdrawal and reinfusion, as compared to the deep capillary layer (Fig.3E-H), even though the latter is more vascularized [21, 22]. OCTA was able to pick up systemic and regional variations in microcirculation when subjected to physiological challenges (Fig.4).

**Fig. 4.**
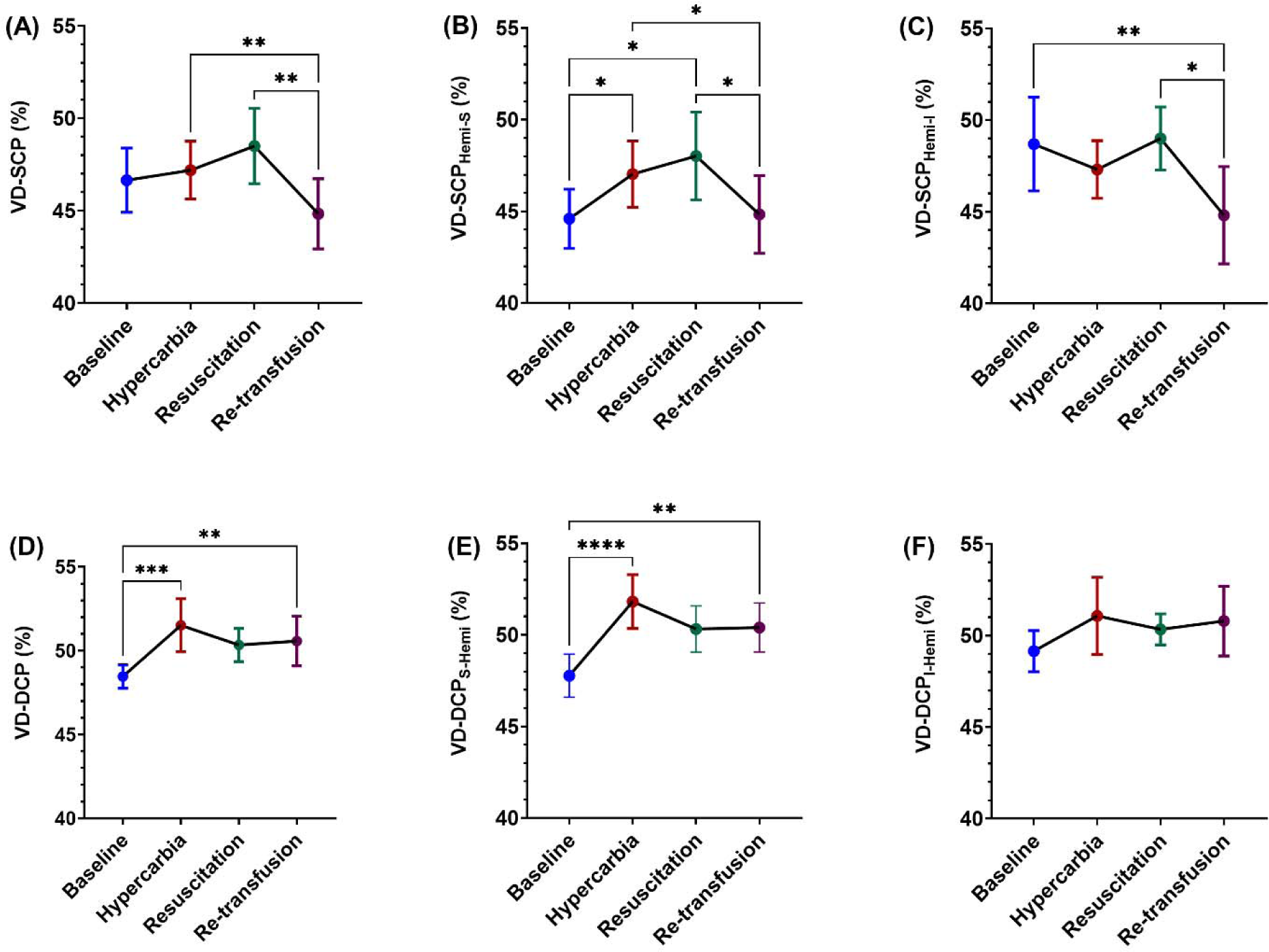
Vascular density parameters (VD-SCP (A), VD-SCP_Hemi-S_ (B), VD-SCP_Hemi-I_(C), VD-DCP (D), VD-DCP_Hemi-S_(E), and VD-DCP_Hemi-I_ (F)) derived from OCTA scan across baseline, hypercarbia, resuscitation, and re-transfusion stages are reported here.

The retina has been long considered as a *window* into the brain with respect to brain and spinal cord related pathologies [23, 24], and we are contributing to establish this eye-brain blood flow connection from a peri-operative perspective in this preclinical OCTA imaging study. Similarities between human and porcine retinal structures have been well established (Table 4) over the years and we extended our understanding of retinal blood flow behavior under experimentally challenging physiological conditions (*hypercarbia*, *resuscitation* by epinephrine, hemorrhage followed by *re-transfusion*, etc.)

In the high CO_2_ condition (*hypercarbia* phase), cerebral and choroidal blood flow increase has been reported in large animals [25–27]; however, retinal blood flow maybe regulated differently in pigs [26]. Retinal blood flow may be intraocular pressure(IOP)-dependent, and IOP increase has been reported in rat models when subjected to acute hypertensive conditions due to respiratory acidosis and hypercarbia [28]. In an acute setting with male Wistar rats, Warner et al. [29] characterized the retinal vascular reactivity to hypercapnia (or hypercarbia) temporally, after inducing experimental aneurysmal subarachnoid hemorrhage. They found that both cerebral and retinal capillary beds showed parallel impairment pattern immediately, and up to 6 hours post-injury, thereby showing that retinal vessels indeed reflect the concurrent impairment of cerebral blood flow.

Newborn piglets exposed to 6 hours of PaCO_2_ of 65 mmHg or 80mmHg exhibited acidosis induced decrease in cerebral energy metabolism, increase in apoptotic genes, alteration of Ca^++^/calmodulin-dependent kinase IV activity, and phosphorylation of cAMP response element binding protein [30], alongside induction of cognitive deficits [31]. We subjected our animals to PaCO_2_ of 70 mmHg for approximately 20-30 minutes; the severity of the experimental condition was therefore close to both studies reported by Fritz and co-authors [30, 31], also confirmed by blood gas analyses (Supplementary information S1).

The deep capillary plexus layer exhibits a denser capillary network than the superficial layer [21, 22] (Table 2). Vascular density of superficial and deep capillary plexus layers showed a 1.16% (non-significant) and 6.31% elevation from baseline, respectively, when subjected to high CO_2_ conditions (Fig. 4). The important regulator of retinal circulation is the diameter of blood vessels and increase in CO_2_ leads to the disruption of transretinal distribution of extracellular H^+^ concentration (pH) [32]. Disruption of H^+^ hemostasis leads to retinal acidosis that eventually affects retinal blood flow and other physiological functions. In addition, hypertension in piglets is known to produce free radicals via the cyclooxygenase pathway, which autoregulates retinal and choroidal blood flow in their ocular tissues [33]. Furthermore, in humans a 6mmHg increase in end tidal PaCO_2_ was shown to increase retinal blood flow by almost 35% [34]. Hypercarbia also induces a shift in lactate levels and could also be a contributing factor to the increased retinal blood flow as well. However, it is still unclear if the cause of increased retinal blood flow is due to pH change, lactate level alteration, or a combination of both factors [32].

As evident from Choi et al. [35] a progressive increase of IOP leads to eventual disappearance of OCTA signals and the retinal flow is not detected after IOP of 60mm Hg in porcine eyes. One possible factor for the disappearance of OCTA signals could be the *collapse* or *compression* of retinal microvasculature due to increase in IOP. However, the relationship between BP and IOP may be complicated due to the inherent autoregulatory mechanisms governing it. Hemorrhage due to hypovolemic shock in our animals led to a reduction in MAP and possibly to a corresponding reduction in retinal flow, which may have influenced adequate signal quality for capturing good quality OCTA images. Although not directly applicable to hemorrhage, a study by Coppolino et.al [36] investigated hourly changes in OCTA of hemodialysis patients during their respective sessions and found little or no changes in flow metrics in this population cohort (other than some structural changes in OCT scans). Zhang et al. showed that an acute increase in human IOP of 10 or 15 mmHg for 2 hours, while the blood pressure is constant, does not alter the OCTA vascular density or nerve head flow biomarkers [37]. It is possible that OCTA image acquisition is suitable for a moderate range of IOP, but extreme IOPs might bring OCTA signal detection to its technical limits.

Conversely, Murphy reported that arterial blood pressure has minimal effect on IOP [38]. We observed that the vascular density of SCP layer exhibits significant, positive association with systolic, diastolic, and mean arterial pressures, respectively (Table 2). Furthermore, our MAP in this investigation ranged from 38.67±4.45 mmHg to 84.83±24.68 mmHg, and for such extreme physiological conditions, we were able to capture OCTA images successfully. If the findings by Choi et al. [35] are accurate, then our experimental stages that did not allow to capture any images (even a trace signal of blood flow in retina) were primarily due to an increase in IOP above 60 mmHg (where OCTA flow signals disappear) and this is extremely high range for a pig (normal IOP for pig 10-25 mmHg). Even though we did not measure IOP in our experiments, it is possible that there is no direct dependence of IOP on blood pressure fluctuations.

As expected, the controlled volume hemorrhage led to a significant reduction in MAP and very likely in retinal blood flow as well. This could explain the paucity of usable images during these stages. Upon restoration of systemic perfusion after re-transfusion, retinal blood flow was partially restored, and OCTA images could be obtained again.

In the SCP layer, average vascular density after re-transfusion was 3.9% lower than the pre-hemorrhage baseline measurements, whereas the post transfusion VD in the DCP layer was slightly higher (∼4.4%) than the baseline measurements (Fig.4). Hence, we propose that (i) there is a direct dependence of systemic blood pressure and retinal blood flow, (ii) autoregulatory mechanisms that control retinal blood flow are influenced by IOP and are only functional within a specific IOP range. Further studies are necessary to understand (i) if clinical scenarios (hypercarbia, hypertension, hemorrhage, re-transfusion, etc.) have a direct impact on IOP; and (ii) if there is an infliction point in systemic blood pressure where OCTA signals cannot be detected.

Taking together, the factors that could potentially affect the retinal blood flow (and by extension - OCTA metrics) are IOP, pH, lactate levels, acidosis, autoregulatory dynamics, choroidal flow and intracranial pressure, [26, 28, 29, 32–35, 37, 39–44]. OCTA was able to detect retina blood flow changes akin to the range of blood pressure changes in the porcine animals. Further studies are necessary to elucidate the exact association of VD with IOP and blood pressure for translation of OCTA technology to the bedside.

### Interpretation of OCTA findings

OCTA imaging of porcine eyes is time-intensive with a low yield of quality images compared to humans subjects. In our case, we faced challenges in high CO_2_ stage, most probably to increased ICP and consecutive ocular pressure and the software mechanisms (especially getting the eyeball to a focal point) did not permit image acquisition for all animals. Along the same lines, when isoflurane concentration was increased to achieve a low blood pressure scenario, the eyeball retracted to the ocular cavity, and it was not possible to take images until isoflurane was lowered to maintenance levels and pupil was visible again. Since the Solix machine is FDA approved for clinical use, it is designed with patients in focus and thus difficult to position a porcine head/snout to accurately place the eyes next to image capture unit.

It is further a well-known fact that a wide range of artifacts are generated in OCTA imaging across manufacturers [45]. We observed several artifacts during our scanning session with anesthetized porcine animals, including projection artifacts [46], which led to a reduction in total scan number and/or exclusion of an entire experimental study timepoint(s). Many clinical OCTA metrics, like the foveal avascular zone (FAZ), are inapplicable due to its absence in pigs (Table 2); therefore, only vascular density (VD) was reported in this study.

Exact interpretation of OCTA data is still a debatable topic amongst ophthalmologists and the substantial heterogeneity within currently reported metrics for retinal perfusion and definition of anatomical layers of retina [4, 47]. With respect to retinal diseases, a deficit of flow of increase or decrease by 30% in a wide-field OCTA is considered meaningful change [48]. However, there is no consensus for OCTA data interpretation in acute blood pressure changes or hemorrhagic conditions. Furthermore, lack of standardized reporting OCTA metrics is another issue since each manufacturer classifies retinal layers differently [47, 49]. Next, region wise variation of VD exists in healthy individuals (not only pathological cases), and device related factors (signal index, etc.), ocular factors (axial length, IOP, etc.), individual factors (age, gender, BMI, etc.) and other factors such as smoking, caffeine, etc., can contribute to OCTA metrics variability as well [12, 43, 50, 51]. We did not post process the obtained OCTA images to obtain VD metrics as every single thresholding or contrast technique leads to a different outcome [52], and there is no reported collective consensus among clinical OCTA users. Table 3 shows a list of large animal studies with OCTA and their associated experimental motivations. SCP vascular density has the least heterogeneity amongst all other metrics derived from OCTA imaging [4] and the most widely used as well. Alnawaiseh et al.’s study [53] with sheep retinal perfusion assessment (SCP only) using OCTA under hemorrhagic conditions reported an average drop of vascular density ∼9% (baseline 44.7% vs hemorrhagic shock 34.5%), and ∼8% gain in vascular density after resuscitation (46.9%). This is the only study that is directly comparable to our efforts and our findings agree with their reported findings that VD of SCP layer increased with increased MAP.

**Table 3.** Detailed comparative account of OCTA literature on large animal models.

### Human-Pig axis

The role of cerebral microcirculatory flow in disease outcomes research remains mostly restricted to lissencephalic murine models that allow genetic manipulation and use of aged models. However, microcirculation in lissencephalic brains may poorly represent gyrencephalic human brains. Thus, porcine studies are necessary, allowing us to not only validate previous murine data [58, 59] but also investigate cerebral microvascular hemodynamics in large complex brains. Animal studies are performed under anesthesia, speculum assisted eye opening, and supine position; this is not a direct replication of the clinical scenario. However, it allowed to manipulate physiology and consecutive OCTA imaging, a compromise necessary for proof of concept. Finally for correct interpretation of retinal images displayed, we highlight the similarities and differences between human and porcine ocular features (Table 4).

**Table 4.** Detailed comparison of human and porcine ocular anatomical features.

Taken together, given the gyrencephalic brain representation and similarities in ocular anatomy, the porcine model is well suited for future investigations of microcirculatory brain dynamics and exploration of new microcirculatory resuscitation management.

### Limitations

The number of animals utilized in this study is low, but the complexity of experimental protocol and OCTA imaging hurdles was a critical bottleneck. Some of the physiological conditions may have resulted in extremely high IOP values that could have resulted in the disappearance of OCTA signals.

### Conclusion

OCTA can detect fast changes in systemic blood pressure and can be utilized for non-invasive cerebral blood flow pattern determination during pre- and post-surgical evaluation of patients.

## Author contributions

Study conception or design – SSP, JMP, UH

Data acquisition – SSP, SI, JSR, KA, AD, MD, UH

Data analysis or interpretation – SSP, SI, JSR, UH

Manuscript writing – All

Funding – UH, JMP

## Acknowledgements

The authors would like to thank Dr. Thomas Floyd and Dr. David Busch for their assistance with the blood gas (iStat) machine.

