## Supplementary material for "A Window into the Brain’s Microcirculation: Retinal Optical Coherence Tomography Angiography (OCTA) as Perioperative Bedside Resuscitation Guide?": Tables 1-4

*Table 1. Experimental stages of the study with detailed interventions are listed here.*

| **Physiological Conditions (clinical scenario)** | **Experimental Intervention** | **Duration** |
| --- | --- | --- |
| *Baseline* | Anesthesia only; no experimental changes | 5 minutes |
| *Hypercarbia* | Hypoventilation (to PaCO_2_ of 70 mmHg) | 20-30 minutes |
| Return to baseline | Adjusted ventilation to PaCO_2_ of 30 mmHg) | 30-40 minutes |
| *Hypotension* | Induced by deep anesthesia (high isoflurane) | 15 minutes |
| *Resuscitation* of hypotension | One dose of Epinephrine (IV bolus) | 1-10 minutes post-injection |
| Return to baseline | Anesthesia only; no experimental changes | 30 minutes |
| *Hemorrhage* | Volume controlled withdrawal via arterial catheter (~800 ml) [9] | 45-60 minutes |
| *Re-transfusion* | Autologous transfusion | 30 minutes |

*Table 2. Correlation of vascular density (SCP and DCP) with physiological parameters. Bold text shows significant correlations between variables (p<0.05).*

|  | **VD-SCP** | | **VD-DCP** | |
| --- | --- | --- | --- | --- |
|  | *r* | *p*-value | *r* | *p*-value |
| *SP (mmHg)* | **0.40** | **0.0176** | -0.12 | 0.5054 |
| *DP (mmHg)* | **0.44** | **0.0081** | -0.15 | 0.3966 |
| *MAP (mmHg)* | **0.43** | **0.0096** | -0.15 | 0.3738 |
| *HR (BPM)* | **0.39** | **0.0207** | 0.16 | 0.5066 |
| *EtCO_2_ (mmHg)* | 0.04 | 0.8194 | -0.13 | 0.4612 |
| *SpO_2_ (%)* | 0.18 | 0.3064 | 0.27 | 0.1201 |

*Table 3. Detailed comparative account of OCTA literature on large animal models.*

| **OCTA Device** | **Species** | **Condition** | **Parameters** | **Relevance** | **Change in quantitative metrics** | **Ref.** |
| --- | --- | --- | --- | --- | --- | --- |
| Solix, Visionix (formerly Optovue), France. | Porcine | Eyes held open with speculum; multiple physiological challenges - high CO_2_, low BP,  epinephrine challenge, hemorrhage, autologous reinfusion | VD | SCP/DCP flow as a biomarker for systemic circulation | VD-SCP showed an average increase in perfusion of ~4% and 1.2% for *resuscitation* and *hypercarbia* stages, respectively  Conversely, *re-transfusion* stage showed an approximate average reduction in vascular density by 3.9% | Our study |
| RTVue XR Avanti, Optovue Inc.,  Fremont, CA, USA | Sheep | Stepwise hemorrhage and resuscitation | VD | Only study that has demonstrated the use of VD as a hemorrhage  Or resuscitation biomarker | ~9% drop in VD due to hemorrhagic shock, and a 7% drop in VD due to resuscitation | [53] |
| Optovue Avanti OCT Angiovue System (Visionix, North Lombard, IL) | Seven 2-month-old female domestic pigs | Anesthetized; Retinal pigment epithelium debridement | Anatomical features due to debridement in choroidal area | Presence of choroidal neovascularization | NA | [54] |
| Spectralis SD-OCT, Heidelberg Engineering, Heidelberg, Germany | Yucanta minipigs (males, 35-45kg) | Specific region of interest was quantified after laser damage | Gray value ratio of laser treated to healthy tissue of choriocapillaris | Laser induced outer retinal and choroidal degeneration | OCTA confirmed laser damage – low or no flow | [55] |
| PLEX Elite 9000, Carl Zeiss Meditec Inc., USA. | Porcine- white Yorkshire pig (N = 4, male and  female, age range: 2–14 months) | Eyes held open with speculum | Perfusion density, VD, fractal dimensions | DVC layer is more vascularized than SVC (approximately SCP layer) in porcine animals | VD magnitude was ~50% VD for porcine subjects | [20] |
| Spectralis OCT2, Heidelberg Engineering, Heidelberg, Germany | Female micro-pigs, 37-59 weeks of age; | Intraocular pressure (IOP) increased in a stepwise manner | Vessel area density (VAD), but larger cilioretinal vessels were removed from analysis | Flow signal ceases after IOP of greater than 60mmHg | SVP and DVP layer VADs reduced by 20% at 50mmHg ocular pressure as compared to baseline (15mmHg) | [35] |
| RTVue XR Avanti OCT system, OptoVue Inc., Fremont, CA, USA | 6 in-vivo rabbit eyes | Evaluate performance of new femtosecond laser system for laser surgery | Structural damage assessment due to laser cut | Anatomical integrity of surgical flaps | NA | [56] |
| Axsun Technology Inc.,  Billerica, MA, USA | Ten isolated pig eyes | Perfused with red blood cells | VD (expressed as number of  Vessel Per 100 mm of Grid Line), length of  visible vessel track, count of visible branch points, vessel track depth, vessel diameter,  angle of vessel descent, and angle of dive | OCTA images were compared to confocal imaging | Smaller vessels were less prominent in OCTA as compared to confocal, especially in deeper retinal layers | [57] |

*Table 4. Detailed comparison of human and porcine ocular anatomical features.*

| **Physiological trait** | **Human** | **Porcine** | **Ref** |
| --- | --- | --- | --- |
| Foveal avascular zone | Present | Absent | Our study; [16, 18-20, 60] |
| Location of retinal vessels | Between inner limiting membrane and inner nuclear layer | Same | [17] |
| Major retinal blood vessels | Deep seated | Superficial in never fiber layer | Our study; [17, 39] |
| Central retinal artery | Single | Multiple retinal vessels entering at the margin of optic disc | [17, 61] |
| Central retinal vein | Single | Single | [17, 61] |
| No. of major retinal arteries and venous branches | 4 | 3-4 | [61] |
| Retinal capillary network | Less distinctly laminar; radial peripapillary plexus is bidimensional | Trilaminar organization with the exception of peripapillary zone (4 layers); 3-4 µm in diameter | [17, 39, 62, 63] |
| Holangiotic type retinal vasculature | Yes | Same | [17, 20, 39, 61, 64] |
| Capillary plexus | Suspended like a hammock between arterioles and venules | Plexus is associated with arterial or venous channels with less evident anastomoses between capillary networks | [17, 62] |
| Retinal arteriovenous crossings | Present | Similar diameters to human; 28.4±3.5 crossings per retina, 23% arterial overcrossings | [65] |
| Radial peripapillary capillaries | Repeated anastomoses with underlying venules | Long straight, undergo repeated anastomoses with underlying arterioles, and very few venous connections | [17, 41] |
| Choriocapillaris | 6.5-9.8 µm in diameter (human eye bank specimens) | 8.9-13.9 µm in diameter (26-33 lbs.; male, Yorkshire pig) | [63, 66] |
| Orbital cavity structure | Bony enclosure | Open; smaller in comparison | [16, 19, 67] |
| Annulus of Zinn | Present | Absent | [16, 19] |
| Retractor bulbi muscle | Absent | Present | [16, 19] |
| Globe length (anteroposterior axis) | 24.15 mm | 24 mm | [16, 68, 69] |
| Circulation of optic nerve head | Present | Similar | [17, 20] |
| Axial length | 17 mm (human infant *in vivo*) | 14mm postmortem (piglets -3–5 days old) ); ~19mm (2–14 months) | [19, 20, 70] |
| Lens diameter | 9 mm (69-70 years old) | 12.5 mm (6-7 months; 225 lbs.) | [71] |
| Lens thickness (maximum) | 4.5 mm (69-70 years old) | 9.2 mm (6-7 months; 225 lbs.) | [71] |
| Choroid thickness | 0.15 mm (69-70 years old) | 0.14 mm (6-7 months; 225 lbs.) | [71] |
| Tapetum lucidum | Absent | Absent | [61, 72, 73] |
| Scleral thickness | 0.3-1.35 mm (69-70 years old) | Comparable; 0.25-90 mm (6-7 months; 225 lbs.) | [71] |
