## Supplemental Table S1 for "A Window into the Brain’s Microcirculation: Retinal Optical Coherence Tomography Angiography (OCTA) as Perioperative Bedside Resuscitation Guide?"

**Supplementary information**

**S1. Blood gas measurements**

Blood gas assessment was performed using CG4+ cartridge on iStat machine (Abbott Inc.). Readings were to confirm the *hypercarbia* stage of the experiment, i.e. high CO_2_ as compared to *baseline*.

Table 1. Blood gas measurements from the experimental study (n=2 animals)

|  | **pH** | **PCO_2_ (mmHg)** | **PO_2_(mmHg)** |
| --- | --- | --- | --- |
| **Baseline** | 7.50 ± 0.04 | 34.65 ± 2.33 | 508.0 ± 56.57 |
| **Hypercarbia** | 7.32 ± 0.05 | 55.75 ± 2.90 | 471.5 ± 116.67 |
